# Psilocybin has no immediate or persistent analgesic effect in acute and chronic mouse pain models

**DOI:** 10.1101/2025.07.06.663398

**Authors:** Nicholas S. Gregory, Tyler E. Girard, Akila Ram, Austen B. Casey, Robert C. Malenka, Vivianne L. Tawfik, Boris D. Heifets

## Abstract

The psychedelic psilocybin may have lasting therapeutic effects for patients with chronic pain syndromes. Some clinical and preclinical data suggest these putative benefits derive from direct analgesic effects. However, this possibility has not been comprehensively tested in preclinical models. Here, we show that psilocybin is not analgesic over a range of doses across multiple pain assays and models of acute and chronic inflammatory, neuropathic, or musculoskeletal pain in mice.

## Introduction

Psilocybin, the prototypical classic psychedelic, produces hallucinogenic activity in humans through activation of discrete 5-hydroxytryptamine receptor (5-HTR) subtypes. A resurgence of interest in the therapeutic potential of psychedelics, particularly psilocybin, has led to testing psilocybin-assisted therapy for a range of neuropsychiatric conditions, most notably depression and substance use disorders^1–4^. Early studies across clinical indications have shown large, rapid and durable therapeutic effects after a single hallucinogenic dose of psilocybin, leading to broad optimism that psilocybin may have a role in other high-impact health conditions such as chronic pain^5^.

Clinical studies performed in the 1960s and 1970s suggested that a hallucinogenic dose of a similar psychedelic, lysergic acid diethylamide (LSD), could reduce pain either by a direct analgesic effect or by globally improving emotional health^5–7^. Exploratory analyses from modern psilocybin trials performed in advanced-stage cancer patients with a variety of pain etiologies have shown improvements in several psychiatric conditions associated with chronic pain, such as anxiety and depression^8–11^, but have noted a heterogeneous analgesic effect of psilocybin^10^. Recent survey studies and pilot clinical trial data suggest that patients with various chronic pain syndromes can effectively manage pain symptoms with psychedelics over a range of doses^11–14^, even suggesting that psychedelics (i.e. LSD) may have direct analgesic effect^15^, although changes in pain beliefs and pain acceptance are also likely to contribute. Modern randomized, placebo-controlled trials for various pain syndromes have yet to be completed.

Recently, a few studies in laboratory animals have suggested that single doses of psilocybin and other psychedelic compounds have potent and lasting analgesic effects in some models of chronic pain^16–19^. However, it is unclear whether these early results generalize across models of chronic pain that are widely used in the field, or whether these findings are species-, dose-, or model-specific.

To address this knowledge gap, we performed *in vivo* experiments to evaluate the antinociceptive (pain-relieving analgesic) activity of psilocybin using a range of doses, in a range of experimental pain models in mice, and in each model performed multiple behavioral tests that assayed different sensory, functional, and affective aspects of pain. We chose a range of well-studied doses of psilocybin (0.3, 2, and 10 mg/kg) that have clear central nervous system activity^20–25^. Further, we tested each of these models, behavioral assays, and doses at acute and chronic time points in both male and female mice. Surprisingly, psilocybin had no analgesic properties in any of these tests, except for cold sensitivity in a neuropathic pain model. However, this result was fully accounted for by psilocybin-induced body temperature dysregulation.

## Methods

### Animals

Male and female (10-16 wk) C57BL/6 wild-type mice were housed 4-5 per cage in sterile ventilated cages by Innovive (San Diego, CA) containing Alpha-dri bedding, Enviro-dri enrichment material (Shepherd Specialty Papers), and *ad libitum* access to pre-filled acidified water bottles (Innovive) and irradiated 18% protein rodent diet (Teklad Global). Animals were maintained on a 12:12 light/dark cycle in an SPF facility and allowed to habituate to the Stanford University Vivarium for at least 1 week prior to testing. All behavior testing took place during the light cycle. All procedures followed animal care guidelines approved by Stanford University’s Administrative Panel on Laboratory Animal Care (APLAC) and the recommendations of the International Association for the Study of Pain.

### Compound administered

Psilocybin was obtained from the NIDA Drug Supply Program and reconstituted to a working concentration of 0.03, 0.2, or 1 mg/kg in sterile saline to keep injected volumes consistent across doses. Morphine sulfate (West-Ward, Tinton Falls, NJ, USA) was obtained at a stock concentration of 8 mg/ml and diluted with saline (0.9%) to obtain a working concentration of 1 mg/ml. Buprenorphine was obtained at a concentration of 0.3 mg/mL and diluted to 0.1 mg/mL working concentration (Par Pharmaceuticals, Chestnut Ridge, NY, USA). Mice were dosed intraperitoneally at 0.3, 2, or 10 mg/kg (psilocybin), 10 mg/kg (morphine), or 1 mg/kg (buprenorphine).

### Head Twitch Response and Rearing Bouts

Before experimentation, animals were acclimatized for at least 90 minutes in a dimly lit room. Subjects were randomized, treated with 0.3, 2, or 10 mg/kg of psilocybin or saline (i.p.), and immediately placed in a transparent acrylic arena (68 x 22 x 26 cm) for observation over 20 minutes. A trained observer (A.B.C.) recorded head twitch responses and rearing behaviors in 2-minute intervals. A head twitch was identified as a rapid, back-and-forth rocking tic movement of the head. Rearing bouts were defined by an unsupported or supported upward movement of the rostral portion of the animal with its forepaws raised off the ground, independent of a grooming bout. After testing, each animal was placed back into its home cage.

### Spared Nerve Injury (SNI) model of neuropathic pain

The SNI model was induced as previous described^26,27^. Briefly, following isoflurane anesthesia (4% induction, 1.5% maintenance in oxygen), the left hind leg was shaved and wiped clean with alcohol and betadine. A 1 cm incision was made in the skin of the upper thigh, approximately where the sciatic nerve trifurcates. The overlying biceps femoris muscles were retracted by blunt dissection, exposing the common peroneal, tibial, and sural branches of the sciatic nerve. Next, the common peroneal and tibial nerves were ligated with 5-0 sutures (Ethilon) and cut, with care not to distend the sural nerve. The skin closed with surgical staples, followed by a Betadine application. During recovery from surgery, mice were placed on a warming pad until awake and achieved normal balanced movement. Mice were then returned to their home cage and closely monitored over the following week for well-being. All behavior tests were performed 7 days post-surgery. Following the SNI procedure, the lateral aspect of the hind paw (innervated by the sural nerve) displays a hypersensitive phenotype.

### Formalin model of neuropathic pain

The formalin model was induced as previously described^28^. Briefly, the mice were anesthetized with isoflurane and the plantar surface of the left hind paw was cleaned with an ethanol wipe. 10 µL of 4% formalin (Sigma-Aldrich) in saline was injected into the plantar surface of the paw using a 30G needle. Animals in the control group received isoflurane anesthesia alone. Acute cold sensitivity was performed 30 minutes after formalin injection. Mechanical and thermal sensitivity experiments were performed starting 24 hr following formalin injection.

### Complete Freund’s Adjuvant (CFA) model of inflammatory pain

The CFA model was induced as previously described^29^. Briefly, the mice were anesthetized with isoflurane and the plantar surface of the left hind paw was cleaned with an ethanol wipe. 10 μl Complete Freund’s Adjuvant (F5881, Sigma-Aldrich) was injected under the epidermis into the plantar surface of the left hind paw using a 30G needle. Animals in the control group received light isoflurane anesthesia alone. All behavior tests were performed starting 24 hr following CFA injection.

### Acidic Saline-Induced Muscle Pain

Acid Saline-Induced Muscle Pain (AIMP) was induced using the repeated acidic saline injection model^30^. Specifically, on day 1 mice were lightly anesthetized with 1–2% isoflurane and 20 µL normal saline adjusted to a pH of 4.0 + 0.1 was injected into the gastrocnemius muscle. This was repeated on day 5. Behavioral testing was then performed starting the next day.

### Laparotomy model of surgical pain

The laparotomy model of surgical pain was induced as previously described^31^. Briefly, following isoflurane anesthesia (4% induction, 1.5% maintenance in oxygen), the abdomen was shaved and wiped clean with alcohol and betadine. Each animal received 1 mL normal saline subcutaneously before a 1 cm incision was made through the skin and muscle at approximately the mid-level of the abdomen. A cotton-tipped applicator was brushed against the peritoneal lining for 30 seconds. Then, a 1 cm portion of the bowel was removed and clamped for 10 seconds before being returned to the abdominal cavity. The muscle layer was then closed with 5-0 sutures (Ethilon) and the skin closed with Vetbond (3M Company). Animals then received i.p. injection of their assigned experimental compound before being allowed to recover for 30 minutes on a warmed pad.

### Mechanical Hypersensitivity

The von Frey filament test for mechanical hypersensitivity was performed as previously described^27^ to assess allodynia in mice. Briefly, mice were placed in plexiglass enclosures mounted onto a testing platform with a metal, perforated floor. Mice were acclimated to the testing chamber for 1–2 days prior to the start of the study (1 h/session). On each testing day, the animals were acclimated in the testing chamber for 1h. Mechanical allodynia was assessed by applying von Frey filaments to the lateral plantar region of each hind paw for approximately 2s per stimulus using calibrated filaments (Touch Test kit, NC12775-99, North Coast Medical). All trials proceeded using an up–down trial design starting with 0.16g filament. A sudden paw withdrawal, flinching, or paw licking was regarded as a nocifensive response. A negative response was followed by the use of a larger filament, while a positive response was followed by the use of a lighter filament.

### Thermal Hypersensitivity

The plantar test was performed as previously described^29^ to assess thermal hypersensitivity. Briefly, the subjects were placed in plexiglass enclosures on top of a glass platform. The animals were acclimatized to the testing environment for 1–2 days prior to the start of the study (1h/session). On each testing day, the animals were acclimatized in the testing environment for 20 min, or until cessation of exploratory behavior. A radiant heat source (Plantar Test Analgesia Meter, Harvard Apparatus) was applied to the plantar surface of the hind paw and the time to a nocifensive response was recorded. A cut-off time of 30 sec was enforced to avoid potential injury due to tissue damage. Two trials were performed on the left (ipsilateral) hind paw to obtain the average reaction time per paw and a third reaction time was obtained if the preceding two values differed by 2 sec or more.

### Muscle Withdrawal Threshold (MWT)

Muscle sensitivity to mechanical stimuli was assessed using muscle withdrawal threshold^32^. Mice were acclimated to the behavior room and restraints in two 5-minute sessions over two days. Testing sessions consisted of three trials, which were averaged for each session. For each trial, the mouse was placed in a gardening glove, and the hind limb was exposed. A force-sensitive tweezer (Bioseb Rodent Pincher – Analgesia Meter, Bioseb, Pinellas Park, FL, USA) was then used to squeeze the belly of the gastrocnemius muscle in a steadily increasing manner such that a peak of 1600 g was achieved over 2 seconds. The trial was terminated when the animal attempted to withdraw the limb or achieved the peak value. Trials that deviated by more than 15% from the targeted rate of force application were discarded and repeated.

### Thermal Place Preference

The affective component of thermal sensitivity was assessed using the thermal place preference test^33^. 24 h after CFA or saline injection, mice were injected with either saline, morphine 10 mg, or psilocybin (0.3, 2, or 10 mg/kg) 60 minutes before testing. Subjects were then placed on an apparatus consisting of adjacent temperature-controlled plates. For heat sensitivity, the plates were set to 30 C (neutral) and 40 C (hot). For cold sensitivity, the plates were set to 30 C and 20 C (cold). Mice were initially placed on the neutral plate and allowed to move freely for 5 minutes while video was recorded. Animal movement was tracked using SLEAP ^34^ and location was scored using SimBA^35^.

### Marble Burying

The marble burying test was performed as described^36^ to measure the effects of pain on repetitive activity in mice. For each trial, an empty standard habitat was filled with 5 cm of flat ⅛” corncob bedding, and twenty-five marbles were evenly spaced in rows in the enclosure. One mouse was then placed into each cage, and left undisturbed for fifteen minutes. Once the mouse was removed, researchers quantified the number of marbles buried (number of marbles that were >⅔ covered in bedding). Marbles are cleaned with dish soap and water between each trial.

### Body Temperature

Body temperature was assessed by recording mice with a thermal camera (e53, FLIR Systems, Goleta, CA). Mice were injected with drug and then immediately placed in a plastic enclosure for recording. The area of the body visible to the camera was tracked using SLEAP^34^. For each image frame, pixel intensities for the area identified as the body were averaged and converted to temperature using the calibration provided by the thermal camera. Frames were then averaged and reported as one-minute bins.

### Mouse Grimace Scale

The Mouse Grimace Scale was assessed using the automated machine learning analysis tool PainFace as previously described^37^. Briefly, mice were placed in a plastic enclosure and recorded for 10 minutes to establish a baseline score. Then, 30 minutes after completing laparotomy and receiving saline or drug injection, mice were recorded

for a second 10 minute session. These videos were then uploaded to the PainFace website (www.painface.net) where they were automatically scored using the default C57BL/6 model. A difference score was calculated by subtracting the baseline score from the post-treatment score.

### Blinding procedures

Behavioral testing and analysis were performed blindly such that the experimenter was unaware of the sham vs. active pain model group assignment, or the drug treatment group.

### Statistical Methods

Cohort sizes were determined based on historical data from our laboratory using a power analysis to provide >80% power to discover 25% differences with p < 0.05 between groups to require a minimum of 4 animals per sex per group for all behavioral outcomes. Data are expressed as the mean +/- SEM. Statistical analysis was performed using GraphPad Prism version 10.1.0 (GraphPad Software). Data were analyzed using One-way analysis of variance (ANOVA), Two-way ANOVA, or an unpaired t-test, as indicated in the figure captions, with post-hoc testing where appropriate. The “n” for each individual experiment is listed in the figure legends.

## Results

### Psilocybin has dose-dependent effects on head twitch and rearing behavior

We confirmed that psilocybin was physiologically active over the dose range (0.3, 2, and 10 mg/kg) by observing a dose-dependent increase in head twitch response (HTR), a proxy for 5-HT_2A_R activity^23,38^ (Fig. 1A), and a decrease in rearing behaviors, a proxy for 5-HT1R activity^39,40^ (Fig. 1B). In both sexes, we observed dose dependent effects of psilocybin (Fig. S1A-D).

**Figure 1.**
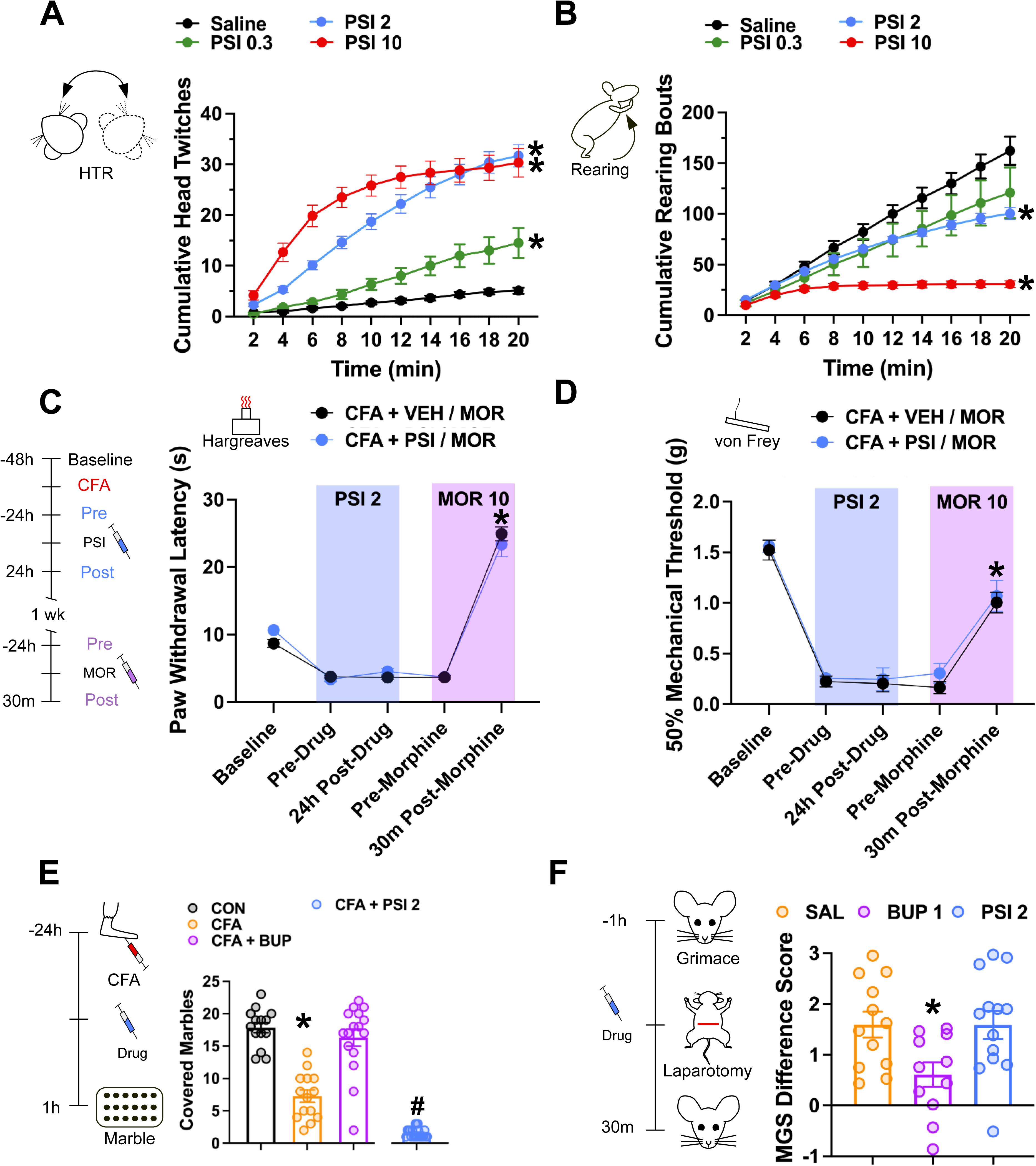
Survey of sensory and analgesic properties of psilocybin. (A) Psilocybin at 0.3, 2, and 10 mg/kg is physiologically active as shown by quantification of Head Twitch Response (males n=3-7 and females n=3 per group). One-way ANOVA of total head twitches F (3, 28) = 42.98 P<0.0001. Post-hoc Tukey Test, * P < 0.05 versus saline. Mean ± SEM. (B) Psilocybin reduces rearing bouts in a dose-dependent manner (males n=3-7 and females n=3 per group). One-way ANOVA of total rearing bouts F (3, 28) = 15.96, P<0.0001. * Post-hoc Tukey Test, P <0.05 versus saline. Mean ± SEM. Psilocybin did not improve mechanical (C) or thermal hypersensitivity (D), whereas morphine produced significant antinociception in both (n=5 males/group). Two-way repeated measures ANOVA (Mechanical: F(2.904, 23.2) = 103.0, P<0.0001; Thermal: F(2.207, 39.72) = 89.85, P<0.0001). Post-hoc Bonferroni test, * P<0.05 across pre- morphine vs 30m post-morphine for mechanical and thermal tests. All data are represented as mean ± SEM. (E) Psilocybin reduces marble burying performance at 2 mg/kg (males n=7-10 and females 7-11 per group). One-way ANOVA of buried marbles, F(5, 90) = 83.07, P<0.0001 P<0.0001. Post-hoc Tukey Test, * P < 0.05 versus control; # P <0.05 versus CFA and control. (F) Psilocybin 2 mg/kg has no effect on mouse grimace scale after laparotomy (males n = 6-7, females = 4-6 per group). One-way ANOVA of mouse grimace scale difference score (post-laparotomy minus baseline mouse grimace scale values), F (2, 33) = 4.535, P=0.0182. Post-hoc Tukey test, * P < 0.05 versus saline and psilocybin.

### Psilocybin has no acute analgesic properties in reflexive, functional, and affective measures of pain in mice

In our initial survey of psilocybin’s sensory and analgesic properties, we tested a dose of psilocybin (2 mg/kg) that has broad effects on acute mouse behavior and immediate early gene expression^20,21,24,41^. At this dose, psilocybin had no effect on the cutaneous mechanical sensitivity of naive mice (Fig. S1E). We then compared the analgesic effects of psilocybin against mu-opioid receptor agonists (morphine 10 mg/kg or buprenorphine 1 mg/kg) in inflammatory and surgical pain models. In the inflammatory model, CFA was injected into the paw, resulting in mechanical and thermal hyperalgesia as well as decreased function on a marble burying task (Fig. 1C-E).

Psilocybin had no effect on mechanical or thermal hyperalgesia (Fig. 1C,D) and further impaired performance on the marble burying task beyond the effect of CFA alone (Fig. 1E). The reduced marble burying effect of psilocybin returned to baseline by 24 hours (Fig. S1F). Consistent with their well-established analgesic properties, morphine significantly reduced mechanical and thermal hyperalgesia, while buprenorphine significantly increased marble burying (Fig. 1C-E). Similarly, psilocybin had no effect on a non-reflexive (“affective”) measure of pain, the mouse grimace scale^37^, measured acutely after a laparotomy. As expected, buprenorphine significantly reduced pain-related facial expressions in this assay^42^ (Fig. 1F). No sex differences were observed in the marble burying task after CFA injection or mouse grimace score after laparotomy (Fig. S1F,G).

### Psilocybin has no immediate or prolonged effects in acute or chronic inflammatory, neuropathic, or centrally-mediated pain models

We next expanded our assessment of psilocybin’s analgesic properties across multiple pain models to include a 3-point dose-response curve at multiple time points (1, 4 and 24 hours after drug injection) during both acute and chronic pain phases (Fig. 2A). We tested multiple sensory modalities in each pain model. As expected, after SNI, mice demonstrated a significant decrease in withdrawal latency when exposed to radiant heat as well as a significant decrease in mechanical threshold, indicating the development of robust thermal (Fig. 2A) and mechanical (Fig. 2B) hypersensitivity at day 7 after injury. None of the psilocybin doses administered altered the established thermal or mechanical hypersensitivity at any of the acute time points tested. Two weeks later, a second dose of psilocybin was given and again, no effect on thermal or mechanical hypersensitivity was observed. We did not observe any sex differences at any time point or dose for SNI (Fig. S2A-D). Together, these findings suggest that psilocybin does not have analgesic properties in a mouse neuropathic pain model.

**Figure 2.**
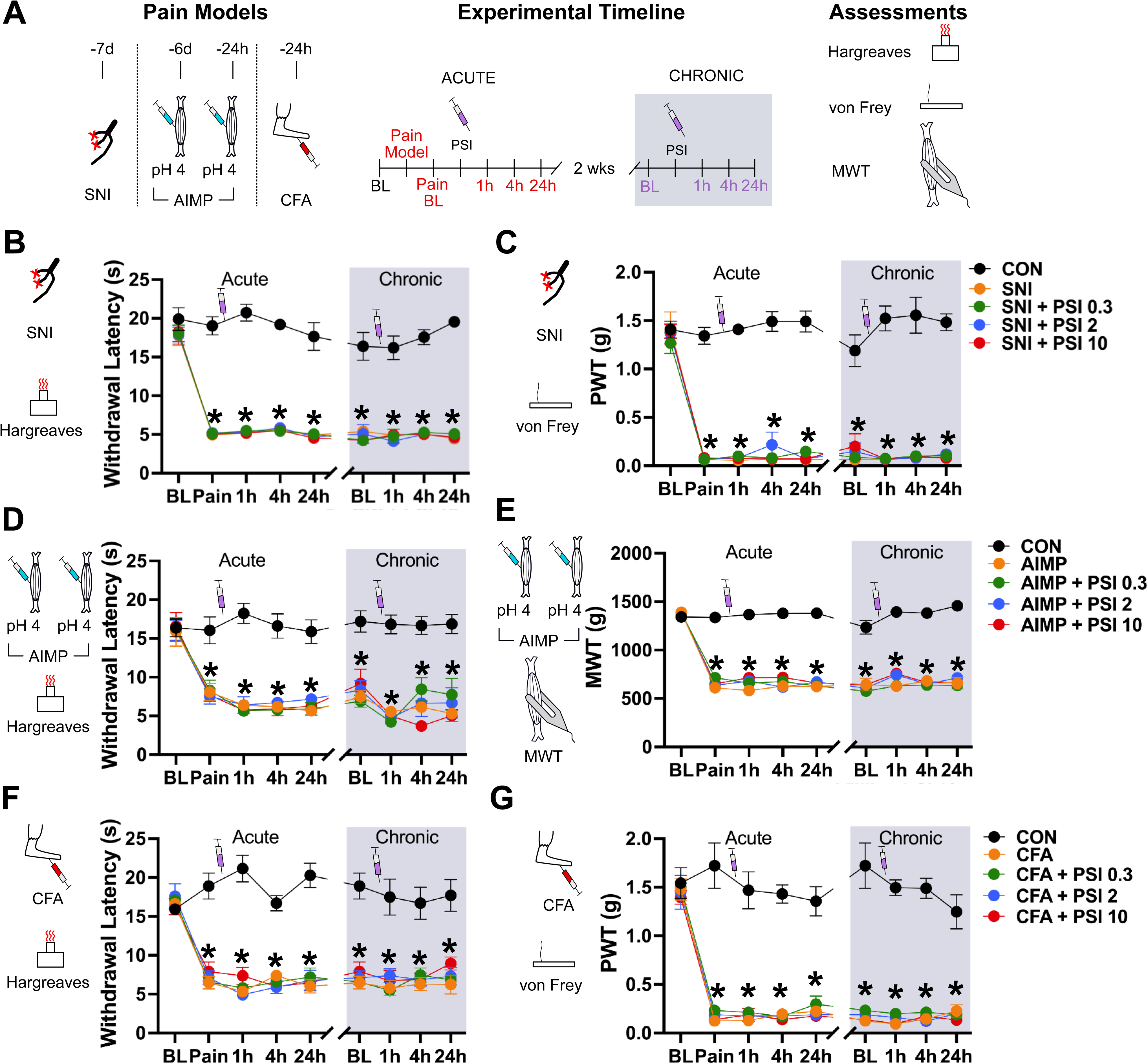
Psilocybin does not improve mechanical or thermal hypersensitivity in male or female mice in three pain models. (A) Legend and timeline for pain models and assessments. Relative to baseline testing: SNI surgery was performed 7 days prior; AIMP involves two acidic normal saline injections, 6 days and 1 day prior; CFA injection to the paw occurs 1 day prior. Psilocybin injections were administered after baseline testing. Hargreaves test (thermal) is quantified by paw withdrawal latency. Von Frey test (mechanical) is quantified by paw withdrawal threshold (PWT). Muscle sensitivity to tweezer force is quantified by muscle withdrawal threshold (MWT). (B,C) SNI produces neuropathic pain and long-lasting thermal and mechanical hyperalgesia. 7 days after surgery, in the acute phase, mice (males and females n=6 per group) were injected with either psilocybin or vehicle control and tested out to 24 hours. Two weeks later, in the chronic pain phase, mice received a second injection and again were tested out to 24 hours. Psilocybin has no effect on either thermal or mechanical hyperalgesia at the doses tested. Two-way repeated measures ANOVA (Thermal: F (32, 440) = 4.861 P<0.0001; Mechanical: F (32, 440) = 7.979 P<0.0001). Post-hoc Tukey Test, * P < 0.0001 versus control. (D,E) AIMP produces long-lasting cutaneous thermal and muscle mechanical hyperalgesia. 24 h after initiation of pain, in the acute phase, mice (males and females n=5-7 per group) were injected with either psilocybin or vehicle control and tested out to 24 hours. Two weeks later, in the chronic pain phase, mice received a second injection and again were tested out to 24 hours. Psilocybin has no effect on either cutaneous thermal or muscle mechanical hyperalgesia at the doses tested. Two-way repeated measures ANOVA (Thermal: F (32, 440) = 2.809 P<0.0001; Mechanical: F (32, 440) = 11.99, P<0.0001) Post-hoc Tukey Test, * P < 0.0001 versus control. (F,G) CFA produces long-lasting thermal and mechanical hyperalgesia due to inflammation of the paw. 24 h after CFA injection, in the acute pain phase, mice (males and females n = 5-7 per group) were injected with either psilocybin or vehicle control and tested out to 24 hours. Two weeks later, in the chronic pain phase, the mice received a second injection and were again tested out to 24 hours. Psilocybin has no effect on either thermal or mechanical hyperalgesia at the doses tested . Two-way repeated measures ANOVA (Thermal: F (32, 280) = 3.664 P<0.0001; Mechanical: (32, 280) = 5.242). Post-hoc Tukey Test, * P < 0.0001 versus control.

We next tested psilocybin on the acid-induced muscle pain (AIMP) mouse model of widespread musculoskeletal pain, which is used to mimic the widespread pain associated with fibromyalgia^32^. This model produced cutaneous thermal (Fig. 2C), muscle pressure (Fig. 2D), and cutaneous mechanical sensitivity (Fig. S2E). None of the psilocybin doses administered improved cutaneous thermal, muscle pressure, or cutaneous mechanical hypersensitivity in the AIMP model in the acute phase (Fig. 2C,D; Fig S2E. Two weeks later, a second dose of psilocybin was given; however, no effect on any of these behavioral measures was observed at any of the time points tested (Fig. 2C,D; Fig S2E). We did not observe any sex differences at any time point or dose for AIMP in tests of mechanical sensitivity of skin or muscle (Fig. S2F-I). We did observe a reduction in thermal sensitivity at a single dose and time point in females (0.3 mg/kg, 4 hours after psilocybin injection at the chronic time point, fig S1K), otherwise psilocybin had no effect on thermal sensitivity. Together, these findings suggest that psilocybin does not have analgesic properties in this model of widespread musculoskeletal pain.

We tested a third model, CFA-induced inflammatory pain. CFA injected under the plantar surface of the hind paw produced thermal (Fig. 2E) and mechanical (Fig. 2F) hyperalgesia of the paw, as evidenced by decreased response latency and reduced paw withdrawal thresholds, respectively. In the acute phase, no dose of psilocybin had an effect on either thermal or mechanical hyperalgesia at any of the time points tested (Fig. 2E,F). Two weeks later, a second dose of psilocybin was administered and, again, we observed no effect at any of the time points tested (Fig. 2E,F). We did not observe any sex differences at any time point or dose for the CFA pain model (Fig S2L-O).

These findings suggest that psilocybin does not have analgesic properties in this mouse model of inflammatory pain.

### Psilocybin drives thermal place preference and cold-insensitivity through hypothermia

We next tested the effect of psilocybin on hot and cold temperature aversion by performing thermal place preference testing in CFA-injected mice. Control mice had no preference for either the 30° C or 40° C surface while CFA-injected mice avoided the 40° C surface (Fig. 3A). Morphine (10 mg/kg, i.p.) abolished this avoidance, while psilocybin-injected mice dose-dependently preferred the 40° C surface (Fig 3A). This preference was observed even in the absence of CFA injection (Fig 3B). Further, psilocybin produced a dose-dependent avoidance of a 20° C surface (Fig 3C), suggesting that the psilocybin does not reduce temperature sensitivity, but rather induces a place preference for warmer temperatures. Similarly, acute psilocybin dose-dependently decreased acute formalin-induced cold sensitivity on the cold-plate test (Fig. 3D). Psilocybin had no significant effect on cold sensitivity in our acute neuropathic pain model (SNI), either at 1h post drug administration (Fig 3E), later post-acute timepoints (Fig. 3F,G), or at any time point after injection in the chronic phase of SNI (Fig 3H-J).

**Figure 3.**
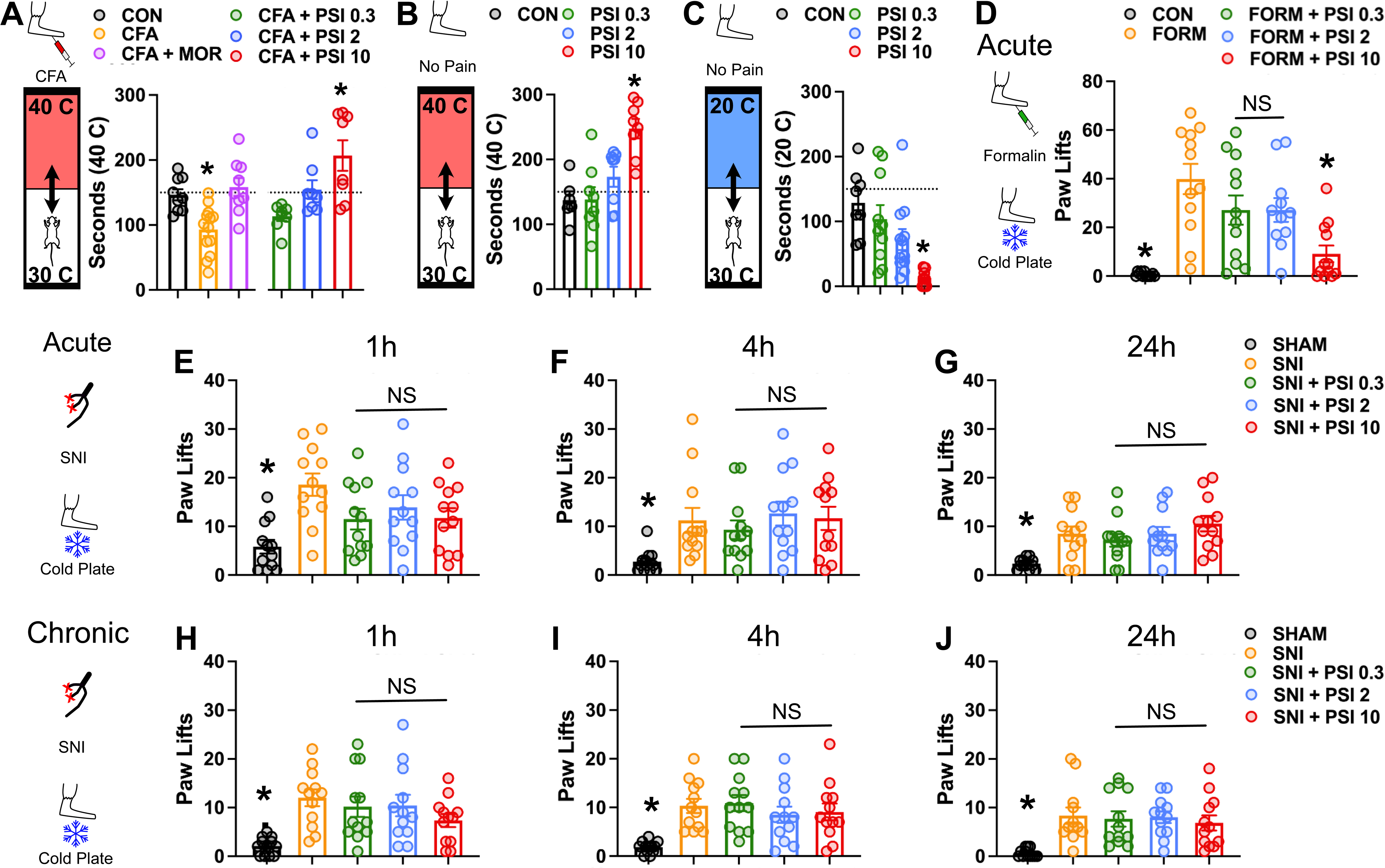
Effects of psilocybin on temperature preference and cold hypersensitivity. (A-C) Thermal Place Preference. Mice are given free access to 30 C and either 40 C or 20 C surfaces. (A) Saline-injected mice have no preference, whereas CFA-injected mice avoid the 40 C plate. This is effect is abolished by 10 mg/kg morphine. Psilocybin produces a dose- dependent preference for the 40 C plate (males n=4-7 and females n=4-6 each per group). One-way ANOVA. F (5, 49) = 9.147 P<0.0001. Post-hoc Tukey Test, * P < 0.05 versus control. (B) The dose-dependent preference is present in mice that are not injected with CFA (males and females, n=3-4 each per group). One-way ANOVA F (3, 26) = 10.34 P=0.0001. Post-hoc Tukey Test, * P < 0.001 versus control. (C) Psilocybin induces a dose-dependent avoidance of 20 C (males and females, n=4-6 each per group). One-way ANOVA F (3, 34) = 7.211, P=0.0007. Post-hoc Tukey Test, * P < 0.001 versus control. (D) Mice (male and female, n =4-6 each per group) were injected with formalin or sham into the paw and given i.p. injection of saline or psilocybin (0.3, 2, 10 mg/kg). 30 minutes later they were assessed for cold sensitivity using the cold plate test. There was a significant decrease in paw lifts compared to SNI controls at 10 mg/kg, but not at 0.3 or 2 mg/kg (F (5, 53) = 8.311, P < 0.0001; Post-hoc Tukey Test * P < 0.05, different from SNI). (E-J) Mice (male and female, n = 6 each per group) underwent SNI and 7 days later were injected with psilocybin or saline control and underwent behavioral testing (acute phase; [E-G]). Two weeks later they received a second injection of the same and underwent repeat behavioral testing (chronic phase; [H-J]). Cold sensitivity was assessed using the cold plate test. In the acute phase (E-G), SNI treated mice had significantly more paw lifts than controls. There was no significant decrease in paw lifts at any dose of psilocybin at the (E) 1 hour, (F) 4 hour, or 24 hour (G) time points (One- way ANOVA, 1 hour: F (4, 55) = 4.840 P=0.0020, 4 hours: F(4, 55) = 5.571, P=0.0008, 24 hours F (4, 55) = 3.520 P=0.0125; Post-hoc Tukey Test * P < 0.05, versus SNI). Two weeks later, in the chronic phase (H-J), psilocybin similarly had no effect at (H) 1 hour, (I) 4 hours, or 24 hours(J) after injection (One-way ANOVA, 1 hour: F (4, 55) = 5.370 P=0.0010, 4 hours: F (4, 55) = 5.571 P=0.0008, 24 hours: F(4, 55) = 6.190, P=0.0003; Post-hoc Tukey Test *, P < 0.05 versus SNI).

Using a thermal camera, we measured the body temperature of mice after injection with either saline or psilocybin and found a dose-dependent decrease in body temperature that emerged at 17 minutes and resolved by 70 minutes, with a nadir around 30 minutes (Fig 4A-D), supporting the hypothesis that observed acute thermal analgesic effects of high-dose psilocybin are best explained by central hypothermia.

**Figure 4.**
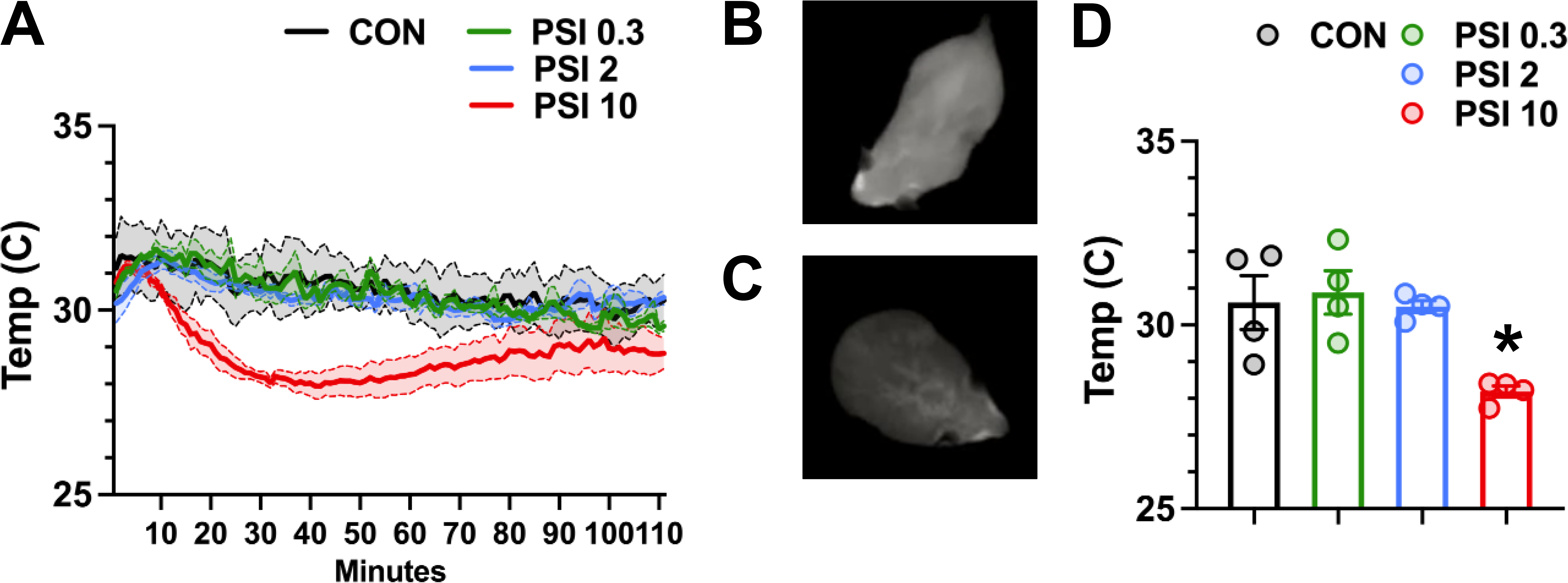
Psilocybin produces dose-dependent hypothermia. (A) Psilocybin produces a dose-dependent decrease in body temperature that emerges at 17 minutes, reaches a nadir around 30 minutes, and normalizes by 70 minutes (male and female mice, n=2 each) Two-way repeated measures ANOVA F (330, 1320) = 2.692 P<0.0001. Post-Hoc Tukey Test (not shown). (B,C) Thermal camera image of female mouse immediately after (B), and 30 minutes after (C) 10 mg/kg psilocybin injection. (D) 30 minutes after injection, body temperature is significantly lower in 10 mg/kg psilocybin as compared to control (males and females, n=2 each per group). One-way ANOVA F (3, 12) = 6.651 P=0.0068. Post-hoc Tukey Test, * P < 0.05, different from control.

## Discussion

The recent interest in psychedelics for the treatment of various psychiatric conditions has prompted speculation that it may also be efficacious in for the treatment of chronic pain. Our experiments are the first to explore the latter possibility in a systematic manner using multiple pain models, both sexes and testing both reflexive and non-reflexive outcomes. Herein we reproduce the finding that psilocybin is physiologically active, in a dose-dependent manner, from 0.3 to 10 mg/kg in HTR and rearing assays; however, we found that none of these doses produced measurable analgesia in any of the acute or chronic pain models tested. In head-to-head comparisons, we show that both morphine and buprenorphine reverse sensory, functional, and affective measures of pain, while psilocybin did not. We then expanded our survey to include both acute and chronic time points, multiple doses of psilocybin, and both sexes in three distinct pain models: spared nerve injury (neuropathic), acid-induced muscle pain (widespread musculoskeletal), or Complete Freund’s Adjuvant (inflammatory). We found that none of the three physiologically active doses possessed any measurable analgesic qualities at any time point in any of these models. We did observe, however, that psilocybin causes a dose-dependent decrease in body temperature, which likely explains the preference for warmer temperature environments in mice treated with psilocybin. This temperature dysregulation may also explain the acute effects of high-dose psilocybin on cold sensitivity.

An important strength of this paper is the diversity of pain models and behavioral tests employed. The tested models have distinct underlying mechanisms providing broad potential to see analgesic effects for a variety of drug classes in current clinical use. Further, the range of behavioral assays used test different features of the pain experience. Reflexive tests depend on sensory transduction at the peripheral tissue which then engages a local reflex arc at the level of the spinal cord. While these tests provide valuable insight into peripheral sensitization and spinal cord mechanisms, they do not necessarily evaluate the unpleasantness or functional impairments caused by a pain state at the level of brain. To address these aspects of pain, we included marble burying as an assay for functional impairment wherein the instinct to bury marbles is opposed by the hyperalgesia triggered by performing the task^36^. We also tested the effect of laparotomy on facial expressions using the PainFace automated scoring tool. Facial expressions reveal the spontaneous, ongoing affective component of pain^37^. This combination of sensory, functional, and affective measures gives a more complete understanding of the pain experience and suggest that psilocybin has no effect on these aspects of pain that can be measured in mice.

Our findings are consistent with existing pre-clinical literature on the complex role of serotonin in nociception and analgesia. Psilocin, the active metabolite of psilocybin, potently activates multiple 5-HTRs, which collectively have varied effects at peripheral nociceptors, spinal cord, and descending pain modulation pathways^43^. Psilocin most notably activates 5-HT_2A_Rs, which mediate the hallucinogenic effect of virtually all serotonergic psychedelics^23,40,44,45^. However, 5-HT_2A_R agonists administered peripherally or intrathecally can worsen pain in animal models, while 5-HT_2A_R antagonists can enhance analgesia^46^. On the other hand, activation of intrathecal 5- HT_1A_R has been linked to stress-induced analgesia through an endocannabinoid-mediated mechanism^47^. Both 5HT_1A_Rs and 5-HT_2A_Rs are expressed in the descending pain modulation system projecting from RVM to spinal cord dorsal horn and have facilitatory and inhibitory effects on pain depending on the cell type and the animal’s conditions^43^. Notably, our results showed no change in mechanical sensitivity in either naïve or injury models, suggesting psilocybin may not affect pain thresholds regardless of this balance. Plasticity-related effects of psilocin and other psychedelics are not clearly associated with analgesia either: trkB activity, ostensibly stimulated by psilocin-induced release of Brain Derived Neurotrophic Factor^44^, is associated with worsened mechanical hypersensitivity^48^ and 5-HT_2A_R-mediated plasticity in the spinal cord may contribute to chronic neuropathic pain^43^.

During our screen for analgesic properties of psilocybin we observed a dose-dependent decrease in body temperature. This finding corroborates previous work ^49^ demonstrating a reduction in core body temperature similar to the time course in our study. Erkizia-Santamaria found a greater magnitude of effect at lower doses, likely due to the greater temperature sensitivity of invasive monitoring. These investigators also provide pharmacologic evidence that 5-HT_2A_R is necessary for the hypothermic effect^49^. Thus, behavioral assays, particularly those that rely on variations in temperature, may be confounded by hypothermic effects of psilocybin. Further, disruption of thermal regulation may be more than an incidental finding. Alitalo et al. demonstrated that drug-induced hypothermia can alter signaling through TrkB, one of the putative mechanisms for psychedelic-induced neuroplasticity^50^. These data suggest that body temperature may be a key parameter that should be monitored and controlled for in future studies of psilocybin.

Our findings contradict recent pre-clinical studies testing various serotonergic psychedelics in other pain models. Kolbmann et al. showed that i.v. administration of psilocin produced immediate and persistent reductions in mechanical and thermal hyperalgesia induced by formalin in rats^16^. This discrepancy may be explained by differences in the route of drug administration or the species used^51^. Lauria et al. found ayahuasca--an herbal brew most commonly made with material from *B. caapi* and *P. viridis* plants containing a range of psychoactive substances, including *N,N*- dimethyltriptamine (DMT) and multiple harmala alkaloids that act on the 5-HT_2A_R--had acute (on-drug) analgesic effects in inflammatory and neuropathic pain models, however, consistent with our findings, no analgesic effects were observed at later time points ^18,52^. These plants also contain monoamine oxidase inhibitors, which alter the metabolism of DMT as well as endogenous monoamines^52^. A recent preprint reports that psilocybin (both 0.3 mg/kg and 1 mg/kg) very modestly (i.e. visible with log-transformation) improved SNI-induced mechanical sensitivity for days to weeks after a single dose. 0.3 mg/kg psilocybin reduced cold sensitivity over a similar time course, but 1 mg/kg psilocybin appeared less effective. The small effect size and inconsistent dose relationship have uncertain biological relevance^19^. Overall, preclinical studies from other groups have not consistently demonstrated post-acute analgesic effects of serotonergic psychedelics, and methodological differences or hypothermic effects may potentially explain differences observed at acute time points.

Human studies of the analgesic properties of psychedelics, however, suggest they may have some beneficial effects on the cognitive-evaluative aspects of pain. Ramaekers et al. found that low-dose lysergic acid diethylamide (LSD) improved cold tolerance and reduced the unpleasantness associated with the cold pressor task ^15^. Notably, this effect was only observed at the highest dose, 20 µg, at which the subjects began reporting psychoactive effects from LSD. This finding is consistent with our pre- clinical data showing an acute reduction in cold sensitivity in mice. Other studies of LSD in cancer patients found the drug improved mood, anxiety, and perceived quality of life^8,9^. These dimensions of pain are outside the scope of rodent models used in our work and merit further evaluation in future studies. Beyond LSD, early-stage trials of psilocybin for headache showed some efficacy; however, the putative analgesic mechanisms for psilocybin may differ from other pain conditions. Specifically, activation of 5-HT_1B_ and 5-HT_1D_ receptors in cerebral vasculature may account for the apparent pain relief in certain headache disorders, rather than changes in pain modulation or neuronal plasticity as previously discussed^53^.

The contrasting results between the present study and data from human subjects highlight important limitations in the use of animal models, particularly mice, in studying of the therapeutic efficacy of psychedelics. While animal models for depression and anxiety exist, other dimensions of the human experience may not be amenable to animal studies. Overall quality of life, meaningfulness of existence, optimism, positive attitudes towards life and oneself, and satisfaction with life are substantially improved by psilocybin in human subjects^8^; however, these characteristics cannot be readily measured in animals. Given the emerging safety data for psilocybin^54^, it may be prudent to focus future efforts in well-controlled human trials where these nuanced effects can be observed. Our data suggest that mice do not reflect direct analgesic effects of psilocybin in either sensory or affective components; should these properties emerge in clinical studies, mouse models may be challenging to reproduce reliably. The potential beneficial effects on anxiety, mood, and motivation may warrant further investigation. Rats or other, higher-order mammals may be more appropriate model systems.

Our study has several important limitations. First, we did not evaluate all clinically relevant pain models, most notably cancer pain, where psilocybin improves mood-related dimensions of pain, or models like complex regional pain syndrome^55^, which may have distinct underlying pain mechanisms. Second, our timeline is limited to approximately 14 days of chronic pain (21 days in the SNI model). It is possible that mechanisms of chronic pain differ across time and changes in the CNS that occur later in chronic pain states may be more amenable to the effects of psilocybin. Third, our experiments only test a limited exposure to psilocybin. While human studies of depression report long lasting effects after a single therapy-supported dose^2^, chronic pain may be more responsive to repeated low dose psychedelics (‘microdosing’), or repeated high dose regimen^56^. Ultimately, the potential applications of psychedelics like psilocybin for the treatment of chronic pain, will require well-controlled clinical trials conducted with sufficient rigor to establish efficacy, and capturing multidimensional outcome measures that reflect the biological, psychological, and social facets of chronic pain.

## ACKNOWLEDGEMENTS

We thank the entire Heifets, Tawfik and Malenka Labs for helpful discussions, and the NIDA Drug Supply Program for supplying psilocybin. This work was supported by a Mentored Research Award from the International Anesthesia Research Society (N.S.G.), NIH grant T32DA035165 (A.R.) NIH grant P50 DA042012 (B.D.H. and R.C.M.), funds from the Department of Anesthesiology, Perioperative & Pain Medicine at Stanford University (V.L.T and B.D.H.) and a grant from the Stanford University Wu Tsai Neurosciences Institute (R.C.M.).

---

**Supplemental Figure 1.**
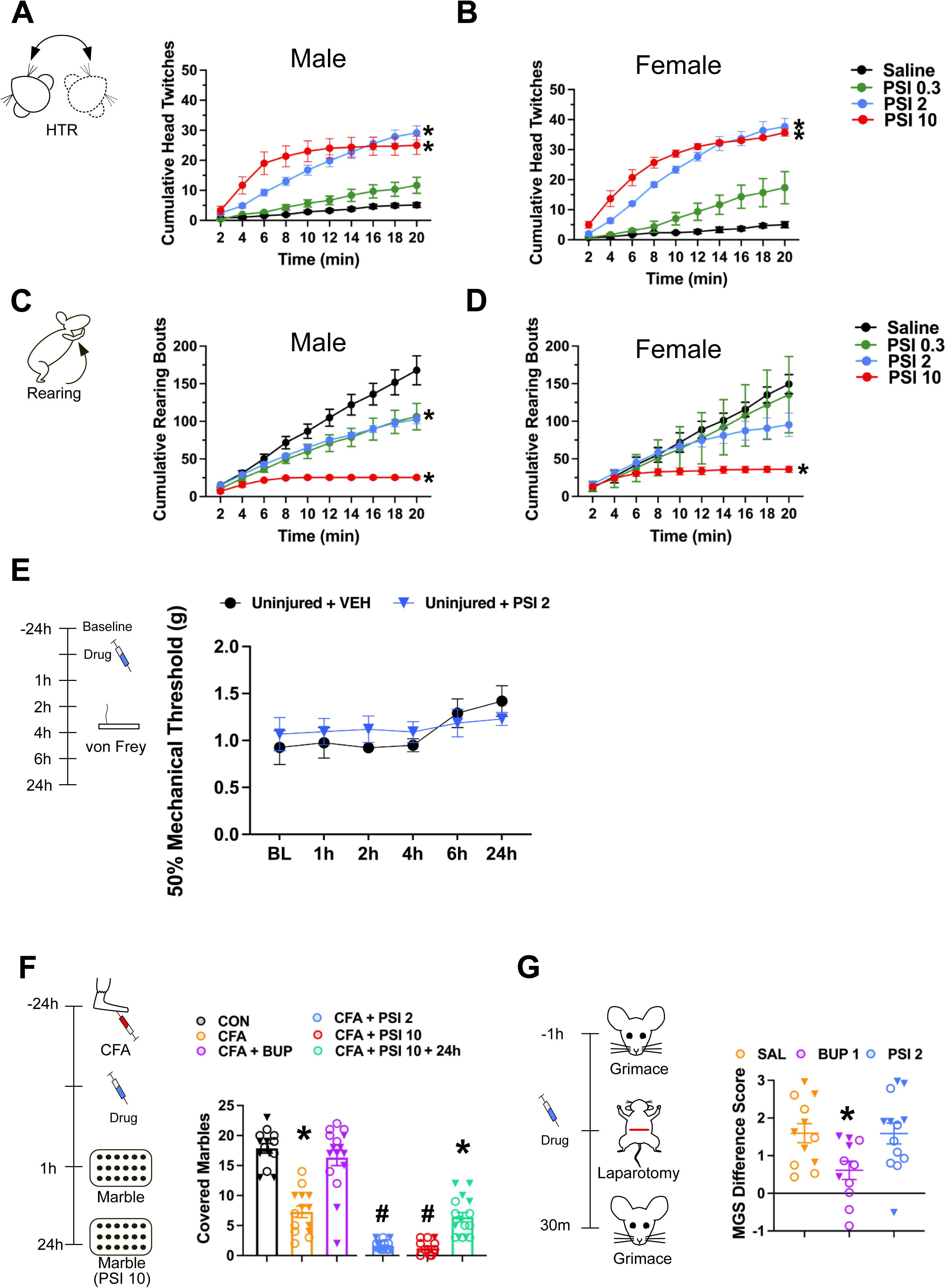
Psilocybin is behaviorally active but does not have intrinsic effects on mechanical sensitivity or affective pain measures in male or female mice. (A,B) Cumulative head twitch response in male (n=3-7) and female (n=3) mice after psilocybin injection. There were significant drug dose and sex effects (Two-way ANOVA, F (3, 24) = 57.23, P<0.0001 for dose, F (1, 24) = 10.94, P=0.003 for sex. Post-hoc Tukey Test, * P < 0.05, versus saline). Males had significantly fewer head twitches at the 2 mg/kg and 10 mg/kg doses. (C,D) Cumulative rearing bouts in male (n=3-7) and female (n=3) mice after psilocybin injection. There was a significant drug dose effect, but no sex differences were observed (Two-way ANOVA, F (3, 24) = 12.84, P<0.0001 for dose, F (1, 24) = 0.05522, P=0.8162 for sex. Post-hoc Tukey Test, * P < 0.05, versus saline). (E) Effect of psilocybin on mechanical sensitivity in uninjured mice. No significant differences were observed between saline control and 2 mg/kg psilocybin at any of the observed time points (Two-way ANOVA, F (5, 42) = 0.6550, P=0.6593) (F) Marble burying in male (n=7-10) and females (7-11) mice after drug injection. There was a significant treatment effect, but no sex differences were observed (Two-way ANOVA, F (5, 83) = 114.7, P<0.0001for treatment, F (1, 83) = 0.0001924, P=0.9890 for sex. Post-hoc Tukey Test, * P < 0.05, versus saline, # P 0.05, versus CFA and saline). Empty circles and solid triangles represent males and females, respectively. (G) Mouse grimace scale difference scores in mice (males n = 6-7, females = 4-6 per group) undergoing laparotomy. Significant drug and sex differences were observed (Two-way ANOVA, F (2, 30) = 3.544, P=0.0415 for drug effect, F (1, 30) = 4.425, P=0.0439, for sex differences. Post-hoc Tukey Test, * P < 0.05 versus saline and psilocyin). Empty circles and solid triangles represent males and females, respectively.

**Supplemental Figure 2.**
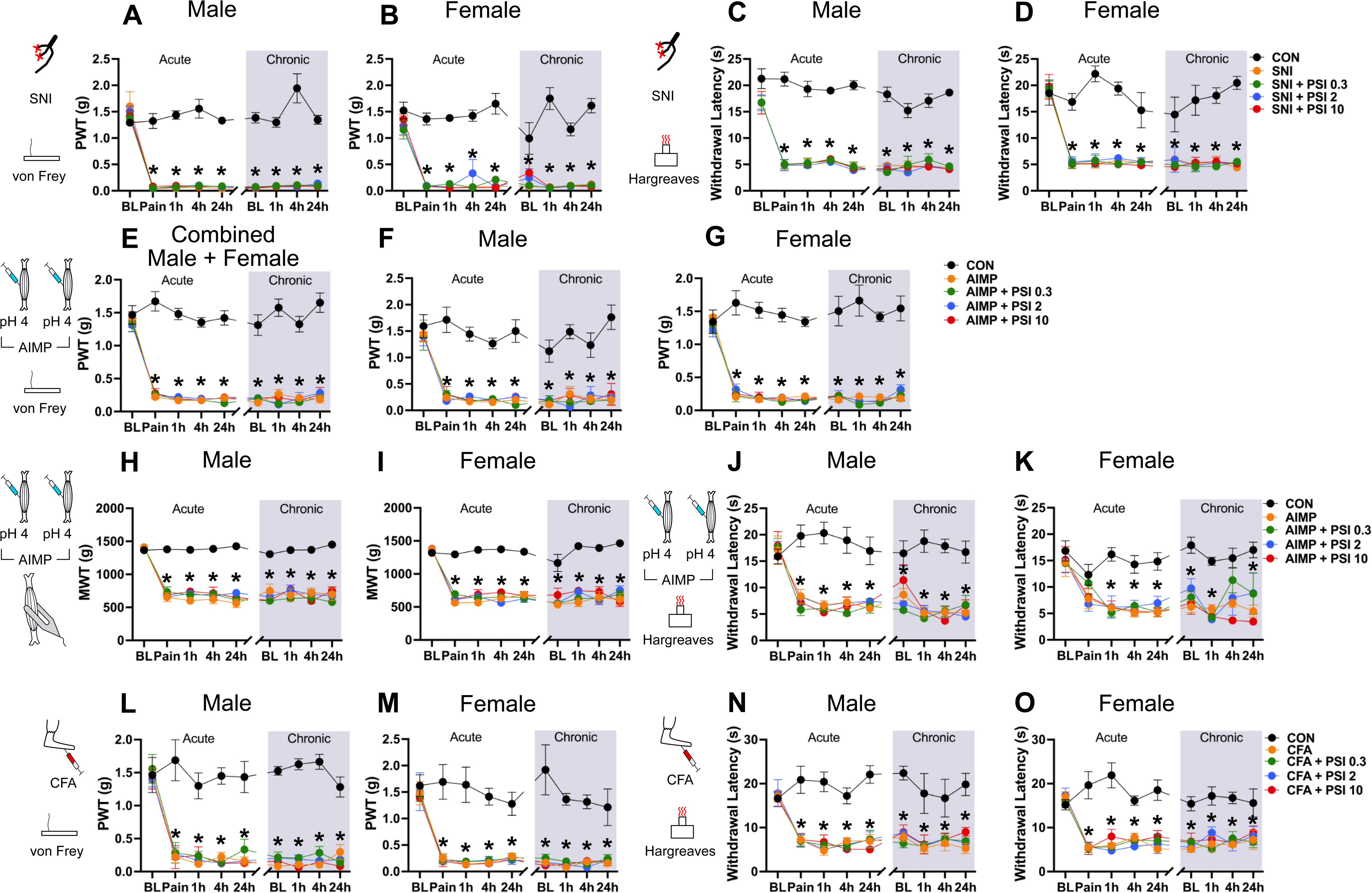
Psilocybin does not improve mechanical or thermal hypersensitivity in male or female mice in three pain models. (A,B) SNI produces similar degrees of mechanical hyperalgesia in (A) male and (B) female mice. PSI had no effect on mechanical hyperalgesia in both male and female mice (Two-way ANOVA, males: F(32, 200) = 9.129, P<0.0001, females F(32, 200) = 3.518, P<0.0001; Post-hoc Tukey Test, * P < 0.05 versus control) (C,D) SNI produces similar degrees of thermal hyperalgesia in (C) male and (D) female mice. PSI had no effect on mechanical hyperalgesia in both male and female mice (Two-way ANOVA, males: F(32, 200) = 2.755, P<0.0001, females: F(32, 200) = 2.956, P<0.0001; Post-hoc Tukey Test, * P < 0.05, different from control). (E) AIMP produces long lasting cutaneous mechanical hyperalgesia 24 h after initiation of pain, in the acute phase, mice (males and females n=5-7 per group) were injected with either psilocybin or vehicle control and tested out to 24 hours. Two weeks later, in the chronic pain phase, mice received a second injection and again were tested out to 24 hours. Psilocybin has no effect on cutaneous mechanical hyperalgesia (Two-way ANOVA, F(32, 440) = 7.979, P<0.0001; Post-hoc Tukey Test, * P 0.05, versus control). (F,G) AIMP produces similar degrees of mechanical cutaneous hyperalgesia in male and female mice. PSI had no effect on mechanical cutaneous hyperalgesia in both (F) male and (G) female mice (Two-way ANOVA, males: F(32, 200) = 2.160, P=0.0007, females: F(32, 200) = 4.456, P<0.0001; Post-hoc Tukey Test, * P < 0.05, versus control. (H,I) AIMP produces similar degrees of mechanical muscle hyperalgesia in male and female mice. PSI had no effect on mechanical muscle hyperalgesia in both (H) male and (I) female mice (Two-way ANOVA, males: F(32, 200) = 6.906, P<0.0001, females: F(32, 200) = 6.164, P<0.0001; Post-hoc Tukey Test, * P < 0.05 versus control). (J,K) AIMP produces similar degree of thermal cutaneous hyperalgesia in male and female mice. PSI had no effect on thermal cutaneous hyperalgesia in (J) male mice (Two-way ANOVA, F(32, 200) = 2.810,P<0.0001). In (K) female mice, PSI treated mice had significantly more thermal cutaneous hyperalgesia than controls (Two-way ANOVA, F(8, 200) = 11.42, P<0.0001), except at a single time point at the 0.3 mg/kg dose in the chronic phase 4 hours after injection, which was not significantly different from control. Post-hoc Tukey test, * P < 0.05 versus control. (L,M) CFA produces similar degrees of mechanical hyperalgesia in male and female mice. PSI had no effect on mechanical cutaneous hyperalgesia in both (L)male and (M) female mice (Two-way ANOVA, males: F(32, 120) = 3.663, P<0.0001, females: F(32, 120) = 1.859, P=0.0088; Post-hoc Tukey test, * P < 0.05 versus control). (N,O) CFA produces similar degrees of thermal hyperalgesia in male and female mice. PSI had no effect on mechanical cutaneous hyperalgesia in both (N)male and (O) female mice (Two-way ANOVA, males: F(32, 120) = 1.674, P=0.0247, females: F (32, 120) = 2.509, P=0.0002; Post-hoc Tukey Test, * P < 0.05).

**Supplemental Figure 3.**
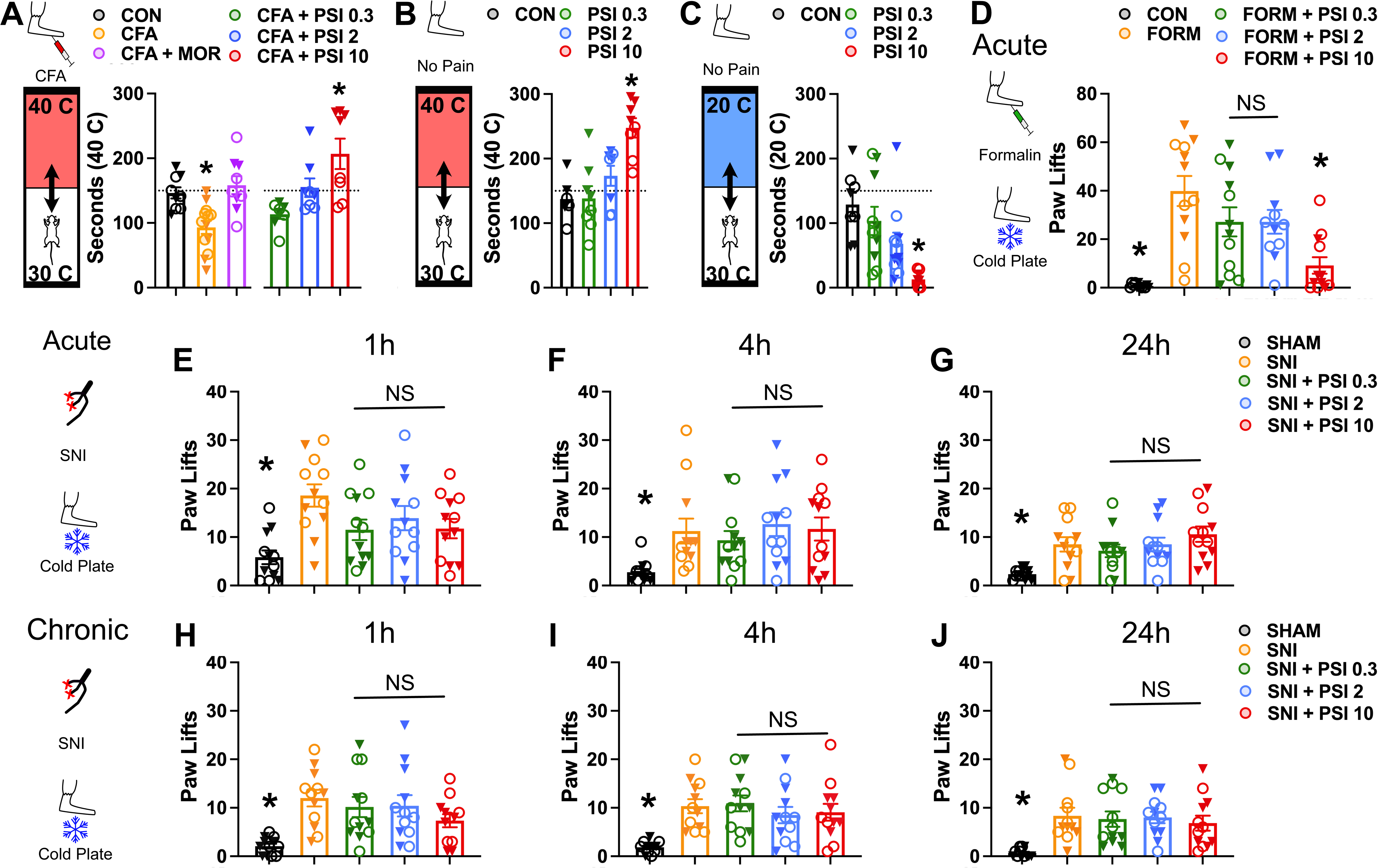
Effects of psilocybin on temperature preference and cold hypersensitivity in male and female mice. Empty circles and solid triangles represent males and females, respectively. (A-C) Thermal Place Preference. Mice are given free access to 30 C and either 40 C or 20 C surfaces. (A) Saline-injected mice have no preference, whereas CFA-injected mice avoid the 40 C plate. This is effect is abolished by 10 mg/kg morphine. Psilocybin produces a dose-dependent preference for the 40 C plate (males n=4-7 and females n=4-6 each per group). Females treated with 2 mg/kg and 10 mg/kg PSI spent significantly more time on the 40C plate than males, otherwise no significant sex differences were observed (Two-way ANOVA, F(5, 43) = 4.784, P=0.0015; Post-hoc Tukey Test, * P < 0.05 versus control). (B) The dose-dependent preference is present in mice that are not injected with CFA (males and females, n=3-4 each per group). No interactions between sex and drug dose were observed (Two-way ANOVA, F(3, 22) = 0.8305, P=0.4913; Post-hoc Tukey Test, * P < 0.05 versus control). (C) Psilocybin induces a dose-dependent avoidance of 20 C (males and females, n=4-6 each per group). No interactions between sex and drug dose were observed (Two-way ANOVA, F(3, 29) = 0.2580, P=0.8550; Post-hoc Tukey Test, * P < 0.05 versus control). (D) Mice (male and female, n =4-6 each per group) were injected with formalin or sham into the paw and given i.p. injection of saline or psilocybin (0.3, 2, 10 mg/kg). 30 minutes later they were assessed for cold sensitivity using the cold plate test. There were no sex differences in cold sensitivity at any of the measured doses (Two-way ANOVA, F (5, 55) = 1.351, P=0.2570; Post-hoc Tukey Test * P < 0.05, different from SNI). (E-J) Mice (male and female, n = 6 each per group) underwent SNI and 7 days later were injected with psilocybin or saline control and underwent behavioral testing (acute phase; [E-G]). Two weeks later they received a second injection of the same and underwent repeat behavioral testing (chronic phase; [H-J]). Cold sensitivity was assessed using the cold plate test. In the acute phase (E-G), SNI treated mice had significantly more paw lifts than controls. There was no significant interaction between sex and drug dose in paw lifts at the (E) 1 hour, (F) 4 hour, or 24 hour (G) time points (Two-way ANOVA, 1 hour: F(4, 50) = 0.6896, P=0.6026, 4 hours: F(4, 50) = 2.211, P=0.0811, 24 hours F(4, 50) = 1.972, P=0.1132; Post-hoc Tukey Test * P < 0.05, versus SNI). Two weeks later, in the chronic phase (H-J), psilocybin similarly had no effect at (H) 1 hour, (I) 4 hours, or 24 hours(J) after injection (Two-way ANOVA, 1 hour: F(4, 50) = 1.972, P=0.1132, 4 hours: F(4, 50) = 0.2676, P=0.8974, 24 hours: F(4, 50) = 0.9760, P=0.4291; Post-hoc Tukey Test *, P < 0.05 versus SNI).

